# Role of aggregate size, multistability and communication in determining cell fate and patterning in *M. xanthus*

**DOI:** 10.1101/627703

**Authors:** Juan A. Arias Del Angel, Natsuko Rivera-Yoshida, Ana E. Escalante, León Patricio Martínez-Castilla, Mariana Benítez

**Affiliations:** Laboratorio Nacional de Ciencias de la Sostenibilidad (LANCIS), Instituto de Ecología, Universidad Nacional Autónoma de México, Mexico City, México; Departamento de Ciencias Naturales, Universidad Autónoma de México, Unidad Cuajimalpa, Mexico City; Centro de Ciencias de la Complejidad, Universidad Nacional Autónoma de México, Mexico City, Mexico; Programa de Doctorado en Ciencias Biomédicas, Universidad Nacional Autónoma de México, Mexico City, Mexico

## Abstract

The emergence of multicellular organisms that exhibit cell differentiation and stereotypic spatial arrangements has been recognized as one of the major transitions in evolution. Myxobacteria have emerged as a useful study model to investigate multicellular evolution and development. Here, we propose a multiscale model that considers cellular adhesion and movement, molecular regulatory networks (MRNs), and cell-to-cell communication to study the emergence of cell fate determination and spatial patterning of *Myxococcus xanthus* fruiting bodies. The model provides a dynamic accounting of the roles of MRN multistability, intercellular communication and conglomerate size in determining cell fate and patterning during *M. xanthus* development. It also suggests that for cell fate determination and patterning to occur, the cell aggregate must surpass a minimum size. The model also allows us to contrast alternative scenarios for the C-signal mechanism and provides stronger support for an indirect effect (as a diffusible molecule) than a direct one (as a membrane protein).

## 2. Introduction

The emergence of multicellular organisms that exhibit cell differentiation and stereotypical spatial arrangements has been recognized as one of the major transitions in evolution (**Maynard-Smith and Szathmáry, 2000**), and it is estimated to have evolved independently about 25 times (**Grosberg and Strathmann, 2007**). While division of labor by cellular differentiation is recognized as a central feature of multicellular organisms, the evolutionary origin of cell fates during the transition to multicellularity remains unclear. Some authors have postulated that multicellular masses appeared first and only later gradually acquired different cell fates and patterns, thus generating spatial differentiation (referred to as “patterning”, **Haeckel, 1874; Arendt, 2008)**. Alternatively, other authors have proposed that even unicellular organisms were capable of differentiation by alternation of cell fates over time, a view derived from a dynamic perspective of development. Thus, as a result of the formation of multicellular masses, organisms spontaneously exhibited the coexistence and patterning of these cell fates. In turn, these cell fates may correspond to stable states enabled by the dynamics of multistable molecular regulatory networks already present in single cells. Under this view, as cells are incorporated into a conglomerate, new local chemical and mechanical microenvironments may bias cells to spontaneously reach different cell fates (**Kauffman, 1969; Furusawa and Kaneko, 2002; Newman et al., 2003; Mora van Cauwelaert et al., 2015**). Importantly, the formation of such microenvironments may require a minimum conglomerate size, and conglomerate size may in turn be constrained by the accumulation of metabolic waste released by the cells (**Asally et al., 2012**) or by mechanical forces acting over the whole conglomerate and the individual cells (**Jacobeen et al., 2018; Rivera-Yoshida et al., 2018**).

In broad terms, multicellular organisms develop through either a clonal (“stay-together”) or aggregative (“come-together”) mechanism (**Tarnita et al., 2013**). While multicellular development and its evolution have been most extensively studied in organisms in which multicellularity is clonal, such as animals and plants (**Grosberg and Strathmann, 2007**), aggregative multicellular organisms are also capable of generating complex structures with different cellular fates and arrangements (**Bonner 1998; Sunderland, 2011; Nanjundiah and Sathe et al., 2011**). However, aside from a few model species (e.g. *Dictyostelium discoideum*; see **Bonner 1998**), the development of aggregative organisms remains largely unexplored. Studying different evolutionary origins and modes of multicellularity will enable comparative analyses that could help to identify both common and lineage-specific aspects in the evolution of cell fate determination and size regulation in multicellular organisms.

Myxobacteria, an order in the delta-proteobacteria, have emerged as a useful study model for investigating multicellular evolution (**Muñoz-Dorado et al., 2016; Arias del Angel et al., 2017**) and elucidating the dynamics behind cell fate determination and patterning in aggregative multicellular organisms. Among myxobacteria, *Myxococcus xanthus* is the most studied species, and transcription factors and signaling pathways involved in its development have been described (**Bretl and Kirby 2016; Kroos 2017; Arias Del Angel et al., 2017, 2018**). When nutrients in the medium are exhausted, vegetative cells (VEG) in *M. xanthus* aggregate into mound-like multicellular structures called fruiting bodies (**Kaiser, 2003)**. During fruiting body development, cells may reach one of three possible cell fates: programmed cellular death (PCD), peripheral cell (ROD) or myxospore (SPO). Inside the fruiting body, cell fates are arranged into two concentric domains: myxospores are concentrated in the inner domain, and peripheral rods in the outer domain (**Sager and Kaiser, 1993a, b; Julien et al., 2000**). Cells undergoing PCD appear to have a broader, although not well characterized, distribution across the fruiting body (**Lux et al., 2004**).

At the intracellular level, *M. xanthus* development initiates with the activation of the so-called stringent response; this mechanism, which is conserved across bacteria, is responsible for genome-wide transcriptional change and survival under stress conditions (**Boutte and Crosson, 2013**), as well as with the sensing of extracellular cues via signaling pathways, such as the A- and C-signals (**Bretl and Kirby, 2016; Kroos, 2017**). The A-signal is a mixture of amino acids and small peptides which freely diffuse in the medium and are thus likely involved in long-range intercellular communication. The C-signal was originally proposed as a membrane protein involved in communication via cell-to-cell contact and has more recently been considered a diffusible molecule (or a producer of them) (**Lobedanz and Løtte-Soggard, 2003; Muñoz Dorado et al., 2016**). These signals are coupled with a complex regulatory network that has been previously shown to be able to reach steady states of the expression profiles of spores, rod and PCD cells (**Arias Del Angel et al., 2018; Supplementary Figure 1**).

In this work, we propose a multiscale model that considers cellular adhesion and movement, molecular regulatory networks (MRNs), and cell-to-cell communication to study the emergence of cell fate determination and spatial patterning of *M. xanthus* fruiting bodies. Specifically, we apply mathematical modeling to gain an understanding of the emergence of spatial arrangements in a population of initially homogeneous vegetative cells whose MRN dynamics are connected through signaling pathways. The model provides a dynamic accounting of the roles of MRN multistability, communication via C-signal and conglomerate size in determining cell fate determination and patterning during *M. xanthus* development. It also suggests that for cell fate determination and patterning to occur, cell aggregates must surpass a minimum size. Finally, the model is employed to contrast the alternative scenarios for the C-signal mechanism, providing support for an indirect effect (as a diffusible molecule), over a direct one (as a membrane protein).

## 3. Methods

### 3.1 Model description

To study cell fate determination and spatial patterning in a virtual population of *M. xanthus* cells, we specified a Glazier-Graner-Hogeweg (GGH) model that considered cellular adhesion and movement, molecular regulatory interactions and cell-to-cell communication (**Figure 1**). In the model, an intracellular MRN specified the internal state of each cell and determined the fate of that cell, as well as the production of intercellular signals. Communication via the intercellular signals mediated the coupling between the individual MRNs; in this way cells could affect the state of MRNs in neighboring cells.

**Figure 1.**
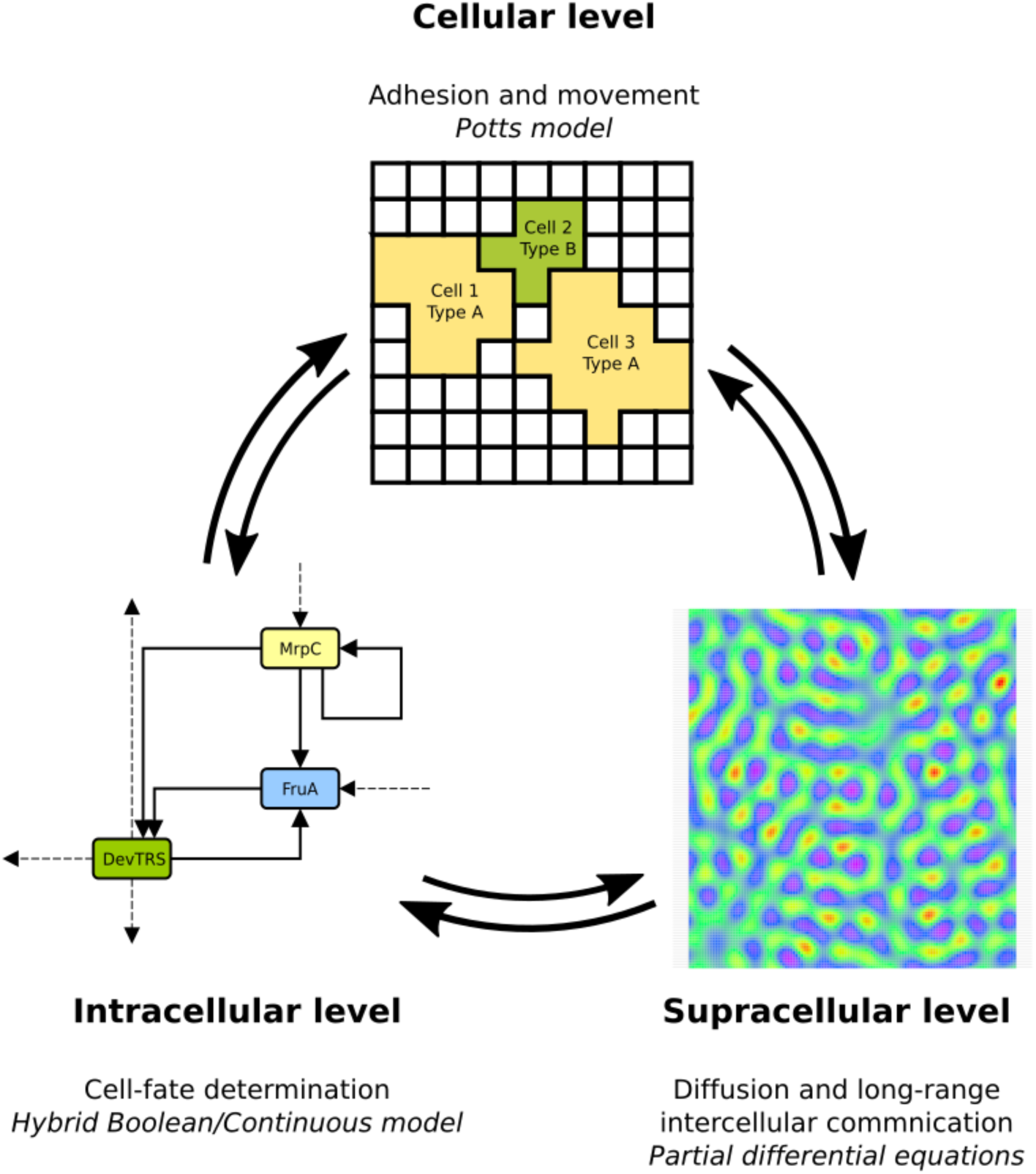
Schematic representation of the Cellular Potts model for cell fate patterning during multicellular development in *M. xanthus*. A Cellular Potts model considering phenomena at the cellular, intracellular and supracellular level is presented. Each level (bold font) captures characteristic phenomena through a different modeling formalism (italic font) and feedback with each other.

The GGH formalism is useful for integrating phenomena at the cellular- and subcellular-levels occurring at different timescales while explicitly considering a spatial domain (**Swat et al., 2012**). In this framework, each scale was captured through a different modeling formalism. The model considered processes at three different spatio-temporal scales, which were all coupled to each other: (1) a dynamic hybrid Boolean/continuous model captured cell fate determination occurring at the intracellular level, (2) the Potts formalism was employed for cell-level behaviors, and (3) partial differential equations were employed for diffusion of chemical fields mediating long-range intercellular communication. Further details of the phenomena and formalisms considered at each scale is presented in the following sections.

#### 3.1.1 Sub-cellular level

At the sub-cellular level, each cell contained an MRN based on experimental evidence that was previously employed to study cell fate determination at the single-cell scale (**Arias Del Angel et al., 2018**). In the MRN, the nodes represented genes, proteins, metabolites or environmental stimuli. At this scale, the MRN dynamics led to multiple steady states that, when compared with the reported experimental data, corresponded to VEG, SPO and PCD. In the MRN, each node could have one of two possible states: 0 if the node was not expressed or was below a certain threshold, and 1 if the node was expressed above the threshold. The state of the nodes changed over time (measured as iterations) according to:

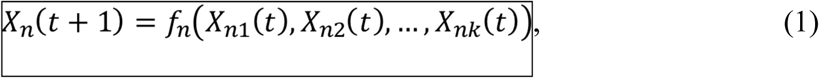

where 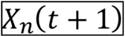 is the state of the node 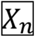 at time 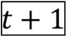 That state was determined by a logical function 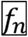, which depends on the state of the regulators 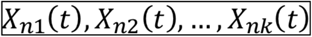 at time 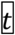. The network was updated using a synchronous scheme such that all of the nodes were updated simultaneously at each iteration.

The MRN presented here was adapted to explicitly consider the role of cell-to-cell interactions. Specifically, the following modifications were made to the previously reported network model (**Arias Del Angel et al., 2018; Supplementary Figure 1)**: (1) the Boolean variables for NUT, ASG and CSG were replaced by continuous variables. This modification facilitated the coupling between cells and the chemical fields while preserving the stepwise behaviour observed in gene regulatory interactions. (2) It was no longer assumed that the state of the transcriptional factor FruA at time (t) (FRUA(t)) was a function of the elements in the C signaling pathway (CSGA(t)) in the same cell. The modified Boolean function for FRUA(t) considered that FRUA(t) was activated when the corresponding cell was surrounded by at least *θ*_CSG_ neighbor cells with CSGA(t) = 1. The complete set of functions specifying the GRN is shown in Supplementary Information 1.

#### 3.1.2. Cellular-level

The Cellular Potts formalism was employed to model cellular adhesion and movement. In this scale, the space was discretized into a regular 1000 × 1000 square lattice with periodic boundaries. Cells consisted of non-overlapping sets of sites over the lattice called pixels. Time was discretized as well into arbitrary units called Monte Carlo Steps (MCS), consisting of a single round of the Metropolis algorithm which allowed the simulation of cell movement across the lattice. The temporal dynamic of the Cellular Potts models is determined by a principle of energy minimization, where energy is specified through a Hamiltonian function 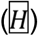 as specified in equation (2).

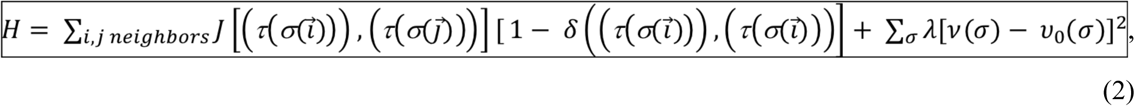

The first and second terms on the right-hand side of equation (2) are the cell-to-cell/cell-to-medium adhesion energies and the volume conservation energy, respectively. The first term on the right side of equation (2), 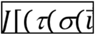))), 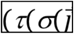)))] is the boundary energy per unit area between two cells 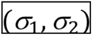 of given fates 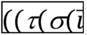))), 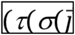))) at a contact point. The term 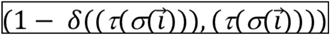 avoids taking into account pixels belonging to the same cell. In the second term on the right side of the equation, 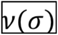 is the actual volume of a cell and 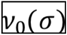 is the target volume. 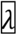 is a constant determining the constraint length.

Two cells were considered to be neighbors if they shared a boundary of at least one-pixel unit. In this model, the values of matrix *J* were considered to be equal over the cell population and thus differential adhesion was not considered. However, values in the matrix *J* representing cell-to-cell (cohesion) interaction strength were allowed to differ from the cell-to-medium (adhesion) interaction strengths. In some versions of the model, the values in the matrix *J* were modified to explore the role of the balance between adhesion and cohesion. The values employed for adhesion and cohesion were 7, 10 and 13 (arbitrary units) in all possible pairwise combinations.

When moving over the lattice, virtual cells attempted to copy their pixel state to neighbor pixels, thus changing *H*. Pixel copy occurred following a Boltzmann probability distribution according to equation (3).

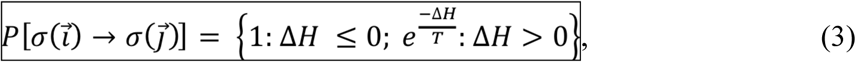

At this scale, cell death was simulated by setting V_0_(σ) equal to zero. This change occurred based on the state of the internal GRN for each cell. For further details about Cellular Potts model specification and implementation see **Swat and collaborators (2012)** and **Glazier and collaborators (2007).**

#### 3.1.3. Supra-cellular level

The supra-cellular level considered diffusion of chemical fields, which mediate long-range intercellular communication. Three chemical fields were considered in the model, representing the available nutrients, A-signal and C-signal. The model assumed that all three chemicals freely diffused across the medium. Also, both A- and C-signal fields were considered to be relatively stable in the medium, with no degradation or consumption occurring outside the cells. Both, A- and C-signal could be freely exchanged between cellular boundaries. In the case of the nutrients, they could be incorporated into the cells, but could not be released back to the medium. Dynamics of the diffusion process for both fields was modeled by partial differential equations as shown in equations (4) and (5).

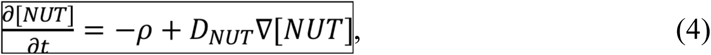

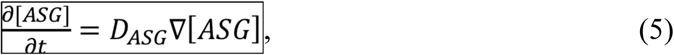

### 3.2 Indexes for assessing spatial organization

To characterize spatial organization, we implemented indexes to quantify the local and global distribution of cell fates in the virtual aggregates. The cellular center of mass, neighborhood and cell fate were recorded for each cell over time. The position of cell fates relative to each other was analyzed for each cell fate by measuring the number of neighbor cells with a given cell fate in order to define the neighborhood preferences, as in **Tosenberger and collaborators, 2017**. The position of the cell fates inside the aggregate was analyzed by calculating the Euclidean distance between the edge of the aggregate (defined as the farthest cellular center of mass from the center of mass of the whole aggregate) and the individual center of mass.

### 3.3. Software and model robustness to parameter variation

To evaluate the sensitivity of the model to parameter variation, we ran the model after modifying the nominal value of key individual parameters. Robustness was assessed by comparing the indexes for spatial organization (section 1.2) obtained for the nominal and modified versions of the model. The tested parameters are highlighted in Table S1.

All models were developed and simulated using CompuCell3D software v.3.7.5 (**Swat et al., 2012**). Because of the stochastic behavior of Glazier-Graner-Hogeweg models, all simulations were repeated at least 30 times. Simulations were visualized using the generic output from CompuCell3D or yEd graph editor v3.16 (**yFiles software, Tubingen, Germany**). The Python library NetworkX v2.1 was employed for network analysis (**Hagberg et al., 2008**) and statistical analysis were performed using R v3.2.3 (**R Core Team 2015**). Graphics were generated using the R package ggplot2 v2.2.1 (**Wickham, 2009**). All models and code applied for analysis are freely available at (https://github.com/laparcela/Myxobacteria-CC3D_model/).

## 4. Results

### 4.1. Coexistence of cell fates is a result of intercellular coupling

We employed the model to study the spatiotemporal dynamics of the developmental process in *M. xanthus*. The system was initialized in the condition representing vegetative growth (VEG). Cells were homogeneously distributed and were allowed to adhere to neighbor cells and to consume nutrients (see Methods and Table S1). As time proceeded, the state of the internal network changed as a response to cell-to-cell interactions and local concentrations of the nutrients and signals. Because the model implemented incorporated the possibility that different cells were exposed to different conditions, the initially homogeneous population segregated into different subpopulations, each one characterized by a different steady state of the MRN (**Figure 2a**) and levels of diffusible elements (**Figure 2b**). In the model, individual cells could reach one of four steady states, associated with cell fates, three of which were also recovered by the single-cell Boolean model previously reported, and correspond to VEG, SPO and PCD cells (**Arias Del Angel et al., 2018**). In the spatiotemporal model, however, an additional steady state was generated and reached by some cells in the population, which matched the ROD profile (ROD cells are characterized by low levels of both ASG and CSG diffusible elements) (**Figure 2a**). This steady state seemed to be generated as a consequence of relaxing the assumption of self-signaling considered in the previous Boolean model (**Arias Del Angel et al., 2018**).

**Figure 2.**
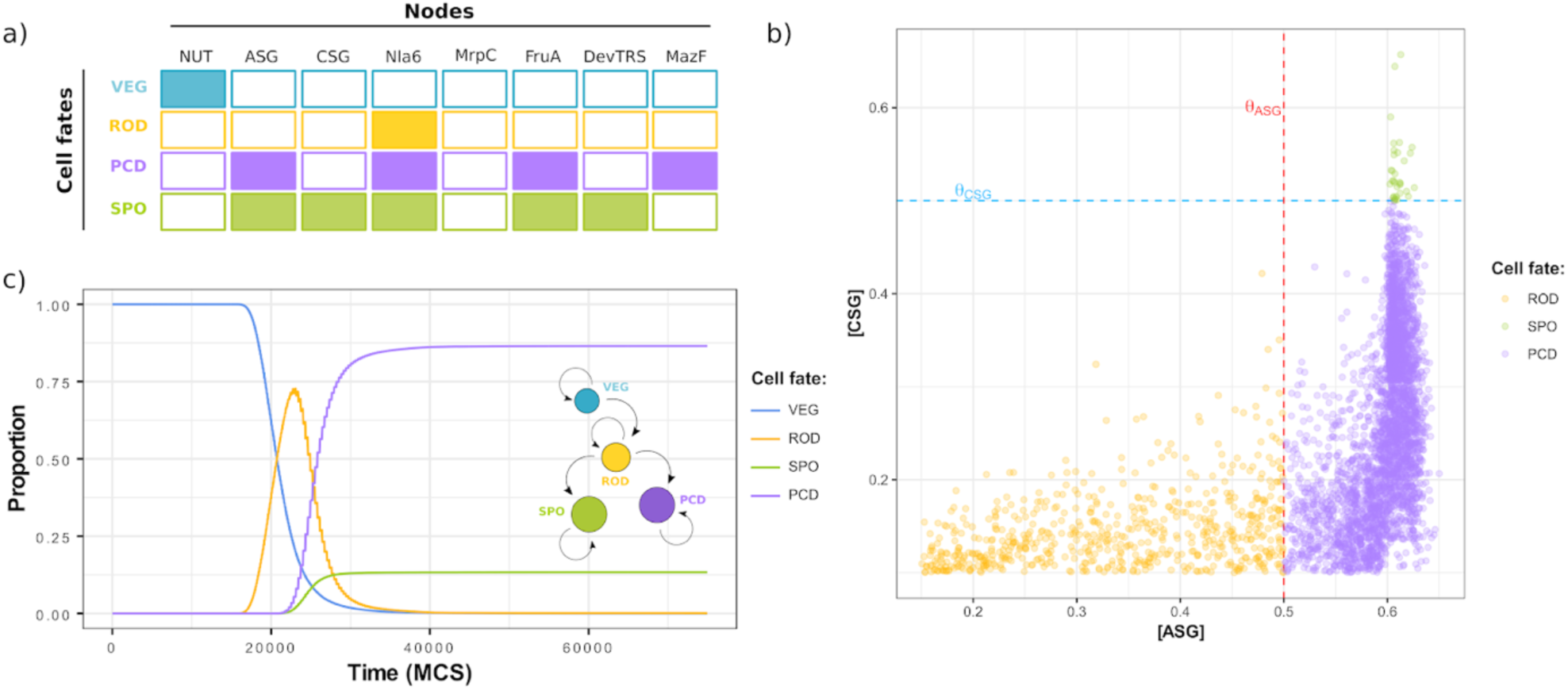
Dynamics for cell fate determination across a population of *M. xanthus* virtual cells. Individual cells in the population reach one of four attractors which are interpreted as cell fates. Cell fates are defined by the state of the internal network **(a)** and relative values of variables representing intercellular communication pathways **(b).** Panel **(a)** shows a subset of the nodes considered in the GRN, which are diagnostic variables that specify cell fates. In **(b)**, dotted lines represent threshold values for ASG and CSG activation. **(c)** Cell fate proportions change over time as a consequence of cell-to-cell interactions. A schematic diagram of the transitions between cell fates is shown inset in the plot (c).

Under a deterministic updating scheme, each attractor recovered in the Boolean model has its own well-defined and non-overlapping attractor basin (i.e., the set of initial conditions leading to the attractor), and transitions between attractor basins were not observed in the absence of stochastic perturbations (**Álvarez-Buylla et al., 2008**). In the spatial model, the local conditions and cell-to-cell interactions to which a cell was exposed enabled them to transit from one attractor to another, causing the proportions of cell fates to vary over time (**Figure 2c**). These transitions were not random, but rather follow an ordered sequence constrained by the dynamics of the internal MRN. Cells remained in the VEG state as long as they had access to nutrients; once nutrients were exhausted, the population rapidly differentiated into ROD cells. ROD cells could remain in this state or differentiate into either SPO or PCD cells (**Figure 2a**,**b**). For cell populations with low NUT, the concentration of A- and C-signals defined a two-dimensional space in which ROD, PCD and SPO can be mapped (**Figure 2b**). In this space, ROD cells were defined by low levels of both A- and C-signals. PCD cells had high levels of A-signal and low levels of C-signal. Finally, SPO cells were defined by high levels of both A-signal and C-signal.

Despite of the stochastic component of the GGH models, the results obtained by the model followed a general trend with low variation across repetitions, indicating that the mechanisms included in the model are sufficient to account for a robust process (**Figure 2c**). This trend held for populations of different sizes above a minimum lattice size of 500×500 (**Figure S2**).

### 4.2. Differentiation and patterning are dependent on the aggregate size

Cell fate determination in individual cells within an aggregate did not occur at random, but rather depended on the local context of each cell, mainly on the interaction with neighboring cells. Moreover, cell differentiation never occurred below a critical aggregate size of ∼50 cells (**Figure 3a**).

**Figure 3.**
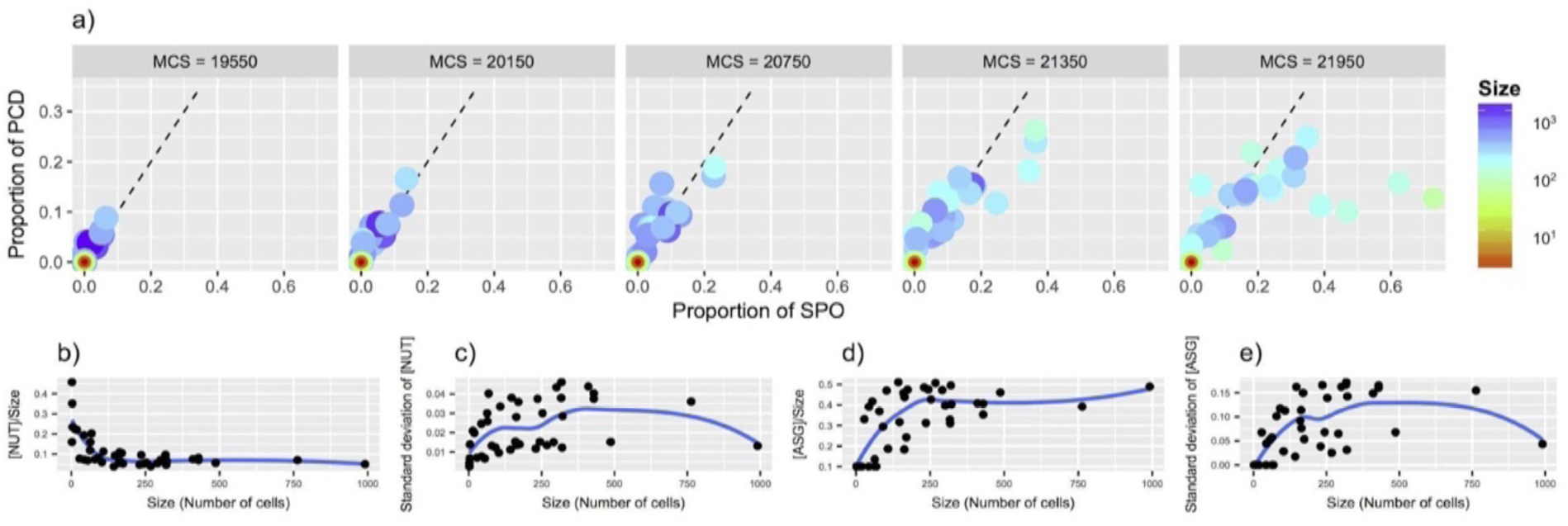
Correlation between cell fate determination and aggregate size. **(a)** Change over time (increasing MCS in panels from left to right) of SPO and PCD proportions at different aggregate sizes. Dotted lines represent equal proportion of SPO and PCD. Colors represent the aggregate size (measured as number of cells in the aggregate). **(b)** Change in the expected level of NUT (nutrient) per cell as a function of aggregate size. **(c)** Change of the standard deviation in NUT (nutrient) levels across the cells inside a single aggregate as a function of aggregate size. **(d)** Change of the expected level of ASG (A-signal) per cell as function of aggregate size. **(e)** Change of the standard deviation in ASG levels across the cells inside a single aggregate as function of aggregate size. In **(b-e)** the blue line represents a polynomial function adjusted to describe the general behaviour of the data.

Since these transitions between cell fates depended on both the depletion of nutrients and the accumulation of A- and C-signals, large cellular aggregates were more likely to reach relatively high concentrations of these signals and locally accumulate them (**Figure 3a**). Small aggregates (and individual cells) accumulated these signals more slowly because a fraction of the signal diffused away into the medium (**Figure 3a and 3c**). Depletion of nutrients and accumulation of A-signal and C-signal did not occur homogeneously within a single aggregate. Also, nutrient and A-signal levels became more heterogeneous among cells as aggregate size increased (**Figures 3c and 3e**).

Within the aggregates, cells at the periphery were more likely to accumulate high levels of nutrients but low levels of A-signal, while the opposite pattern was observed for cells at the interior of the aggregates (**Figure 4**). Since local concentration of the nutrients and signals biased the state of the internal network, their gradients might generate positional information that trigger cell fate determination. The emerging cell fates displayed a preferential position relative to the center of mass of the aggregates and to the cell fates in their neighborhood (**Figure 5**). This cell fate patterning, as well as the change in proportion of cell fate subpopulations over time (**Figure 1c**), was robust to the specific numerical values of key parameters included in the model. Nevertheless, cell fate determination, proportion and patterning were differentially sensitive to specific parameters (**Figure S3 and S4**). For instance, variations in the diffusion rate of the A- and C-signals seemed to affect the cell fate proportions, but not the presence of all four cell fates.

**Figure 4.**
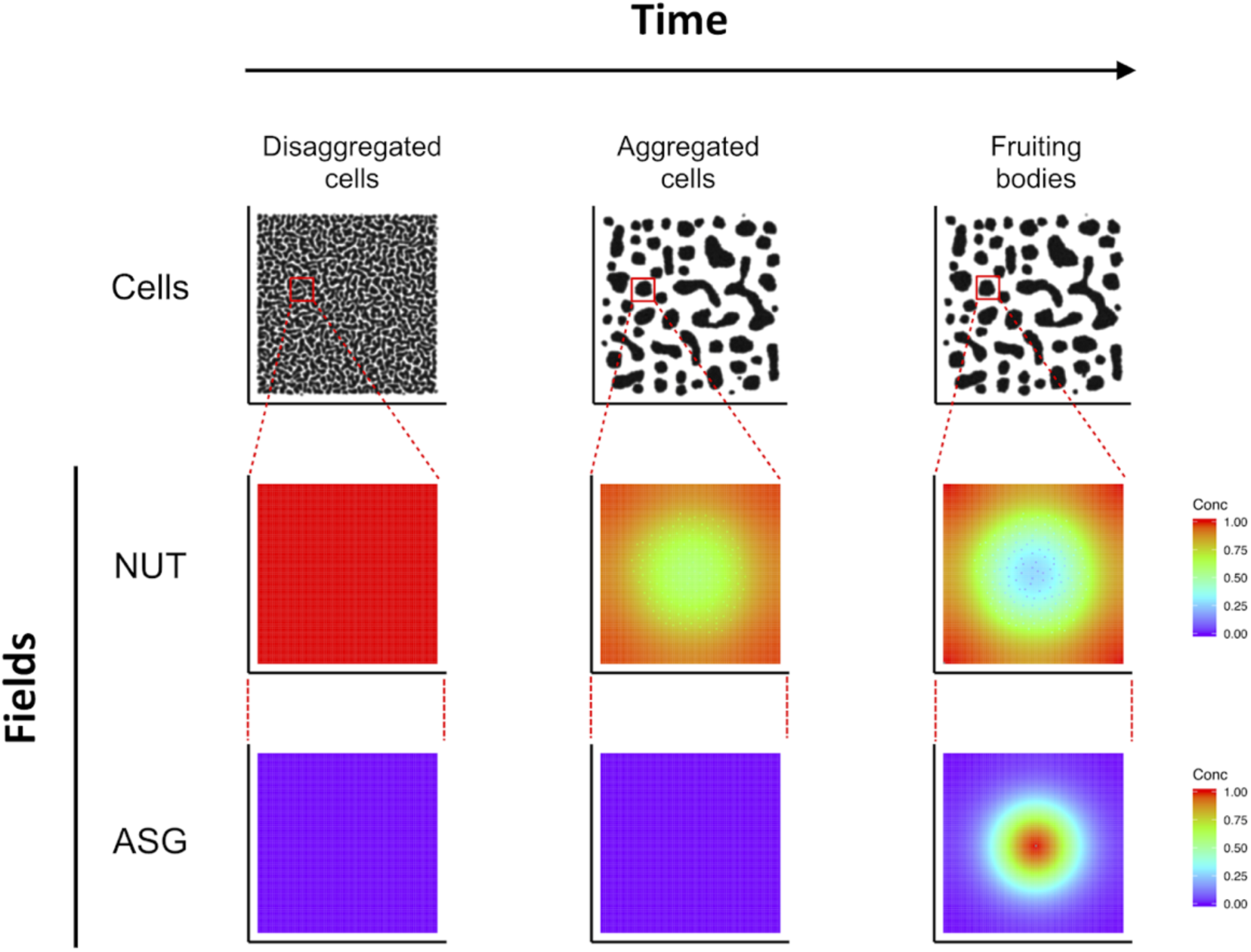
Spatial dynamics recovered by the model for *M. xanthus* FB development. Each row represents a different level and each column a representative time point for the main events recovered by the model (movement of disaggregated cells, aggregation of the individual cells and differentiated fruiting bodies). For the NUT (nutrient) and ASG (A-signal) gradients, only the portions delimited by the red circle in the cellular field are shown and amplified.

**Figure 5.**
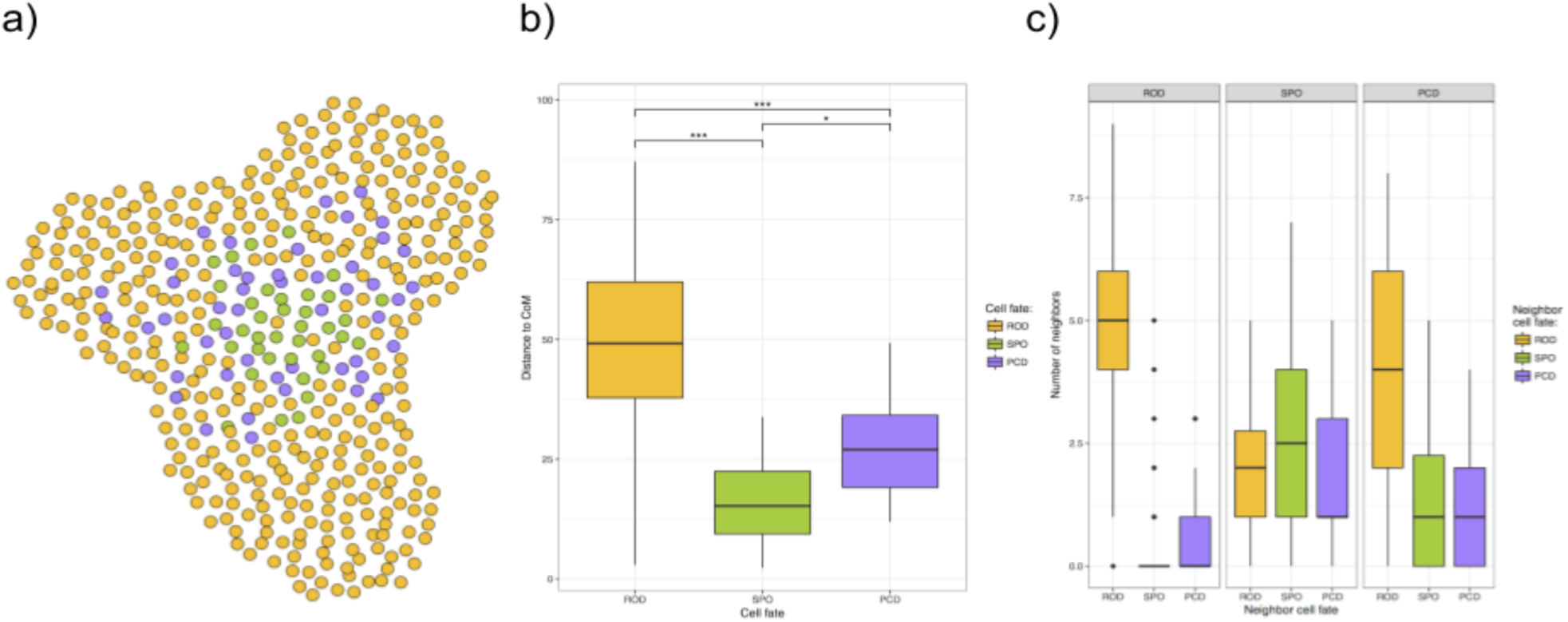
Spatial patterning of cell fates within aggregates. **(a)** Illustrative figure of the spatial distribution of cell fates in a virtual aggregate. Only aggregates of ≥100 cells were included. Cells are displayed as circles representing the position of the center of mass for each cell. **(b)** Distribution of distance of cell fates from the center of mass (CoM) of each aggregate. ANOVA p-values: (*) p < 0.05, (***) p < 0.001 **(c)** Distribution of cell fate of the neighbors of each of the cell fates observed for each aggregate.

### 4.3. Balance between cohesion and adhesion determine aggregate size distribution

The aggregate size distribution was the result of both the strength of cell-to-medium (adhesion) and cell-to-cell (cohesion) affinities. This balance is known as wettability and can be specified at the cellular level through the Potts formalism (Swat et al., 2012). When cohesion was stronger than adhesion, cells tended to form larger aggregates, and vice versa. Different values of wettability resulted in qualitatively different distributions of the aggregate sizes, with a non-linear response to changes in either adhesion or cohesion (**Figure 6**).

**Figure 6.**
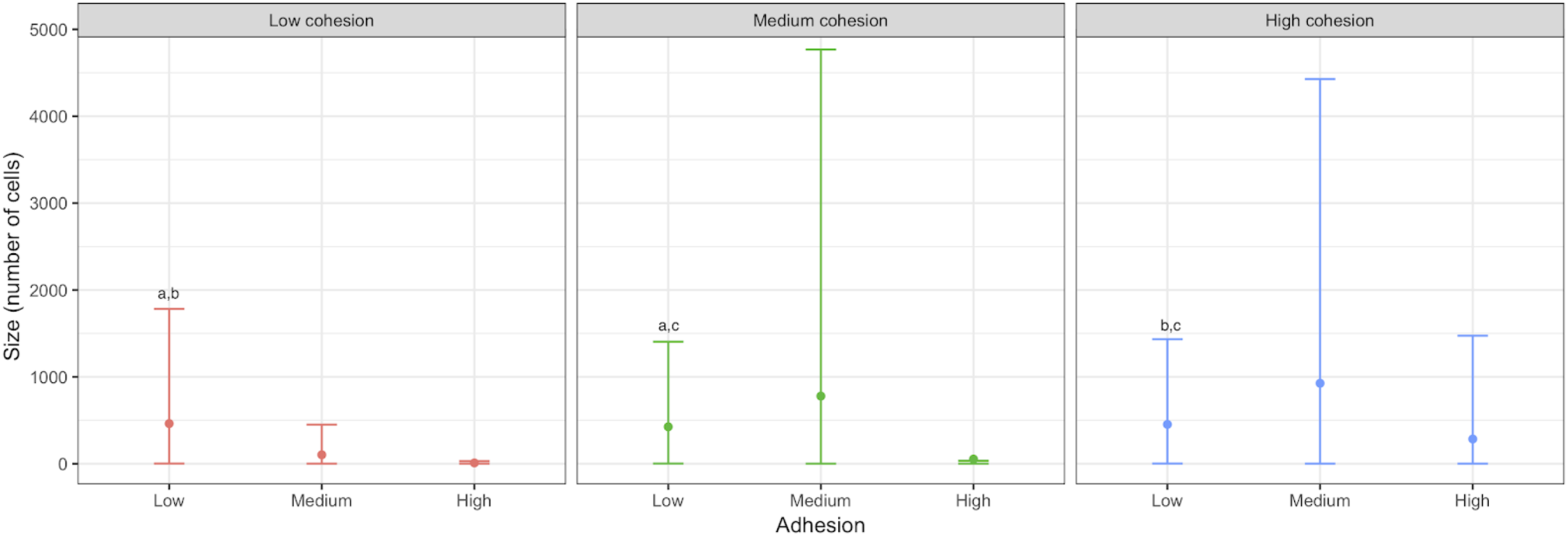
Theoretical effect of the balance between cell-to-medium (adhesion) and cell-to-cell (cohesion) contact energy in aggregate size. For each combination of adhesion and cohesion energy contact, the mean aggregate size (point) and standard deviation (error bars) are shown. Aggregate sizes were measured once the model reached equilibrium. Kolmogrov-Smirnov test to compare distributions; Indexes a, b, and c indicate *p* > 0.05 for pair-wise comparisons. Only non-significantly different pairs of distributions are indicated. *p-values* are adjusted using Holm-Bonferroni method for multiple comparisons.

### 4.4. Contrasting scenarios for the molecular nature of the C-signal

Once we were confident that the model could reproduce the key features of cell fate determination and patterning in *M. xanthus*, we employed it to test the plausibility of two contrasting scenarios. C-signal was initially suggested to be a membrane protein mediating direct cell-to-cell communication (**Lobedanz and Løtte-Soggard**), but recent evidence proposes that it is (or gives rise to) a diffusible molecule involved in indirect cell-to-cell communication. We developed two versions of the model that capture these two alternative scenarios. We found that while both versions of the model were capable of recovering most of the tested properties, only the model considering C-signal as a diffusible molecule was able to recover the appearance of isolated spores (i.e., outside of multicellular aggregates) that has been previously reported (**Higgs et al., 2014**) (**Table 1)**.

**Table 1.** Comparison between alternative scenarios of CsgA-mediated intercellular signaling. using the two models involving CsgA key properties.

| Property | Model 1. Csg-direct | Model 2. Csg-indirect |
| --- | --- | --- |
| *Description* | CsgA acts as an intercellular protein via direct cell-to-cell contact (Løtte-Soggard *et al.*, 2001) | CsgA diffuses in the medium and mediates short-range intercellular communication (Muñoz-Dorado 2016; Robeltzki *et al.*, 2008) |
| *Cell fate trajectories* | Matches experimental evidence | Matches experimental evidence |
| *Predicted cell fate proportions* | SPO < ROD < PCD | SPO < ROD < PCD |
| *Mutant phenotypes* | Matches experimental evidence | Matches experimental evidence |
| ***Isolated myxospore formation (independent of FB development)*** | **Not predicted by the model** | **Predicted by the model, in agreement with previous reports (Yang and Higgs, 2014)** |

## 5. Discussion

Multicellular development is characterized by cell differentiation and patterning. For *Myxococcus xanthus*, our study system, a previous network model grounded on experimental evidence, accounts only for single-cell processes (**Arias Del Angel et al., 2018**), and it does not capture events at the population or multicellular scales at which multicellular development and spatial patterning occur. In this work, we present results of a dynamic multiscale model based on the Cellular Potts formalism to explicitly consider cell movement, gene regulatory interactions and intercellular communication among individual cells during fruiting body formation in *M. xanthus*. Specifically, our results support a dynamic accounting for the origin of cell fates in the transition to multicellularity, in which coexisting cell fates may arise with the aggregation of cells from the coupling of multistable networks already present in single cells (**Kauffman, 1969; Furusawa and Kaneko, 2002; Newman et al., 2003; Mora van Cauwelaert et al., 2015**). This contrasts with a vision in which multicellular masses appear first, and then gradually acquire different cell fates and spatial differentiation **(Haeckel, 1874; Arendt 2008)**.

When the model is employed to simulate fruiting body development in response to starvation, individual cells aggregate, and the population transits from the state representing vegetative growth to those representing the cell fates that emerge during fruiting body. Even when the population consists of initially homogeneous cells, the differentiated local environment inside the aggregates allow them to reach different steady states corresponding to cell fates. It is worth mentioning that in this spatiotemporal model, an additional steady state representing the peripheral rods, emerged as the result of the interaction between cells and their medium. An outcome of the model is that it predicts the stereotypic sequence of VEG -> ROD -> SPO or PCD cell fate determination that emerges from the heterogeneous accumulation of nutrients and signals and the MRN dynamic (**Figure 2**). These results suggest that during multicellular development in *M. xanthus*, the coupling among cells may be important in generating the whole spectrum of cell fates. Given the genetic homogeneity of the cell population, this supports the idea that cellular differentiation can occur in a multicellular context in the absence of genetic variation (**Turing 1952; Von Dassow et al., 2001; Benítez et al., 2008; Mora van Cauwelaert et al., 2015**). Hence, the change of scale during aggregation may on its own constitute an important source of developmental innovation, and changes in the cell-to-cell interactions may render developmental variation.

In addition to cell fate determination, positional information is also generated as a consequence of the formation of gradients of nutrients, A- and C-signals, which translate into cellular patterning with non-random preferential positions independent of any pre-pattern. The cell pattern exhibited qualitative robustness against the specific parameter values employed in the model. The position of SPO relative to ROD recovered by the model is in agreement with experimental evidence that has shown that fruiting bodies are patterned into two concentric domains, with myxospores preferentially located in the inner domain and peripheral rods in the outer domain (**Sager & Kaiser, 1993a, b; Julien et al., 2000**). To the best of our knowledge, the position of PCD cells has not been clearly determined experimentally, but evidence from **Lux and co-workers (2004)** suggests that they occur broadly over the entire fruiting body. Our model suggests that PCD tend to occur in an intermediate ring (**Figure 4**). Because of the limited evidence, this result provides a prediction for future experimental work.

The proposed model suggests that a threshold aggregate size must be surpassed to trigger cell fate determination and patterning. In relatively small aggregates, cells are unable to reach high enough levels of diffusible substances (A- and C-signal) because a significant portion of these signals is lost to the medium through diffusion. In large aggregates, self-activation feedback loops compensate for the proportion of the signals lost to cell-to-medium diffusion and intracellular levels of these signals reach high enough values to trigger downstream effects in the MRNs. In fact, previous studies have considered conglomerate size as a key factor favoring and constraining complexity during development and evolution of multicellular organisms (**Bonner, 1998a**). Moreover, other authors have argued that as cells are incorporated into a conglomerate, new local chemical and mechanical microenvironments may give rise to cues biasing cells toward certain steady states (**Furusawa & Kaneko, 2002**). Overall, our results support the idea that aggregate size may be a cue for development on the basis of a data-grounded model for *M. xanthus* as a model organism.

As previously mentioned, the nature of C-signal has been recently discussed in light of new experimental research. Specifically, the C-signal was originally suggested to be a membrane protein capable of mediating direct intercellular communication (**Lobedanz and Søggard-Anderson, 2003**). In contrast, recent evidence supports an alternative scenario where the C-signal is instead a diffusible molecule (or perhaps involved in the synthesis of such a molecule), mediating indirect cell-to-cell communication (**Muñoz Dorado et al., 2016**). Here, we employed the dynamic model to test the plausibility of these two contrasting scenarios and found more support for the scenario of a diffusible signal mediating indirect cell-to-cell communication. The proposal of a diffusible element is also supported by bioinformatic analyses suggesting that *csgA*, the gene encoding for C-signal, does not contain any putative membrane-anchoring sequence (**Lee et al., 1995**).

Additionally, our model provides insights into the role of MazF, the only reporter marker for PCD. Evidence for the role of MazF has been controversial because its effect has been validated only in a strain where its action might be mediated by interactions with the specific genetic background (**Boynton et al., 2013; Müller et al., 2013**). (**Nariya & Inouye, 2008; Lee et al., 2012; Boynton et al., 2013**). We were aware of these limitations when we included this element in the model; we conclude that, regardless of the specific molecular identify of MazF, PCD-inducing factors acting in other strains may be under direct or indirect regulation of MrpC, as assumed in our model for MazF. Overall, the model as presented renders precise predictions that may ultimately inform experimental work.

In summary, we provide a dynamic accounting of cell fate determination and patterning in *M. xanthus*, which generates several testable predictions. As ongoing research reveals further details of the developmental mechanisms in other myxobacteria, models like the one proposed here may enable comparative studies of developmental processes and dynamics (**Nahmad et al., 2008; Arias Del Angel et al., 2017; Benítez et al., 2018**), thus shedding light on the generic and particular aspects of different fates and instances of multicellularity.

## 6. Acknowledgements

Juan A. Arias Del Angel is a doctoral student from Programa de Doctorado en Ciencias Biomédicas, Universidad Nacional Autónoma de México and was supported by CONACYT 619809. Natsuko Rivera-Yoshida is a doctoral student from Programa de Doctorado en Ciencias Biomédicas, Universidad Nacional Autónoma de México and was supported by CONACYT 580236. The authors acknowledge support and access to computational resources provided by the office of computing at Laboratorio Nacional de Ciencias de la Sostenibilidad (LANCIS-UNAM), particularly from the head of the department, Rodrigo García. The authors thank the CompuCell3D (CC3D) team for workshops, help and feedback about model implementation and software usage, as well as Emilio Mora Van Cauwelaert, Mariana Esther Martínez-Sánchez, Mónica L. García-Gómez and Alessio Franci for useful discussion, and Karen Carrasco-Espinosa, Alejandra Hernández-Terán, Marcelo Navarro-Díaz and Karla Peña-Sanabria for feedback during manuscript preparation. M. Benítez was funded by UNAM-DGAPA-PAPIIT (IA200714, IN113013-3) and CONACyT (221341), and A. E. Escalante by the University of Minnesota, CONACyT postdoctoral fellowship (126166), and PAPIIT-UNAM (IA200814).

## Supplementary Figures

**Figure S1.**
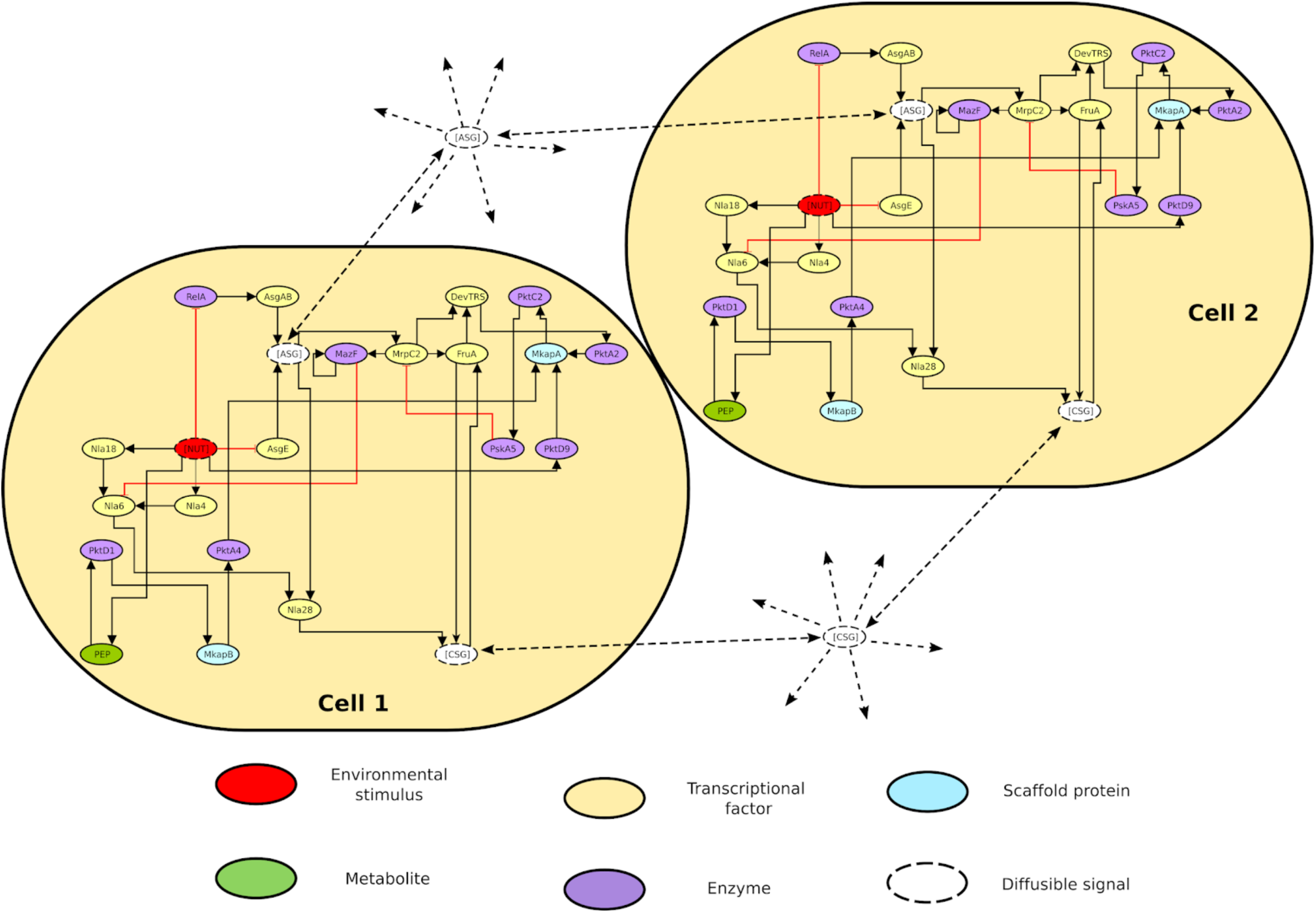
Schematic representation of the internal molecular regulatory network. Two cells are represented (each containing an identical schematic of the molecular regulatory network). Solid lines represent intracellular regulatory interactions. Black and red arrows stand for positive and negative regulatory interactions, respectively. Dotted lines indicates diffusion. Nutrients, A-signal and C-signal are abbreviated as NUT, ASG and CSG, respectively. All other nodes are annotated as usually found in literature.

**Figure S2.**
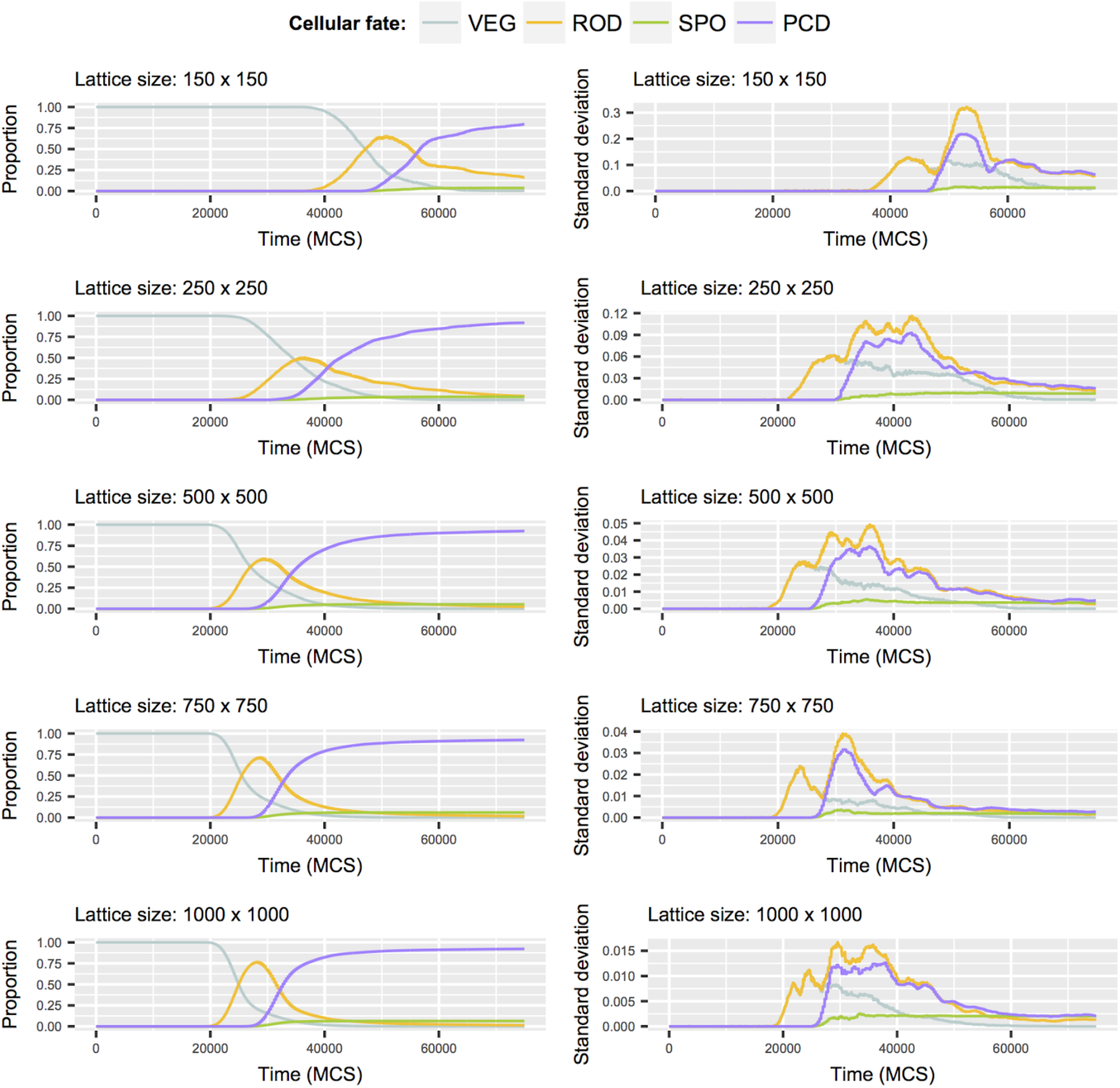
Robustness of system dynamics for large lattice sizes. Each row represents the results obtained for the cell fate trajectory over time when different lattices sizes are used. The left column shows the cell fate trajectories over time (line shows mean of N = 30 simulations). The right column shows the standard deviation for cell fate trajectories obtained from N = 30 simulations. Note that for small lattice sizes, trajectories are delayed and noisier (larger standard deviation) compared to larger lattices sizes.

**Figure S3.**
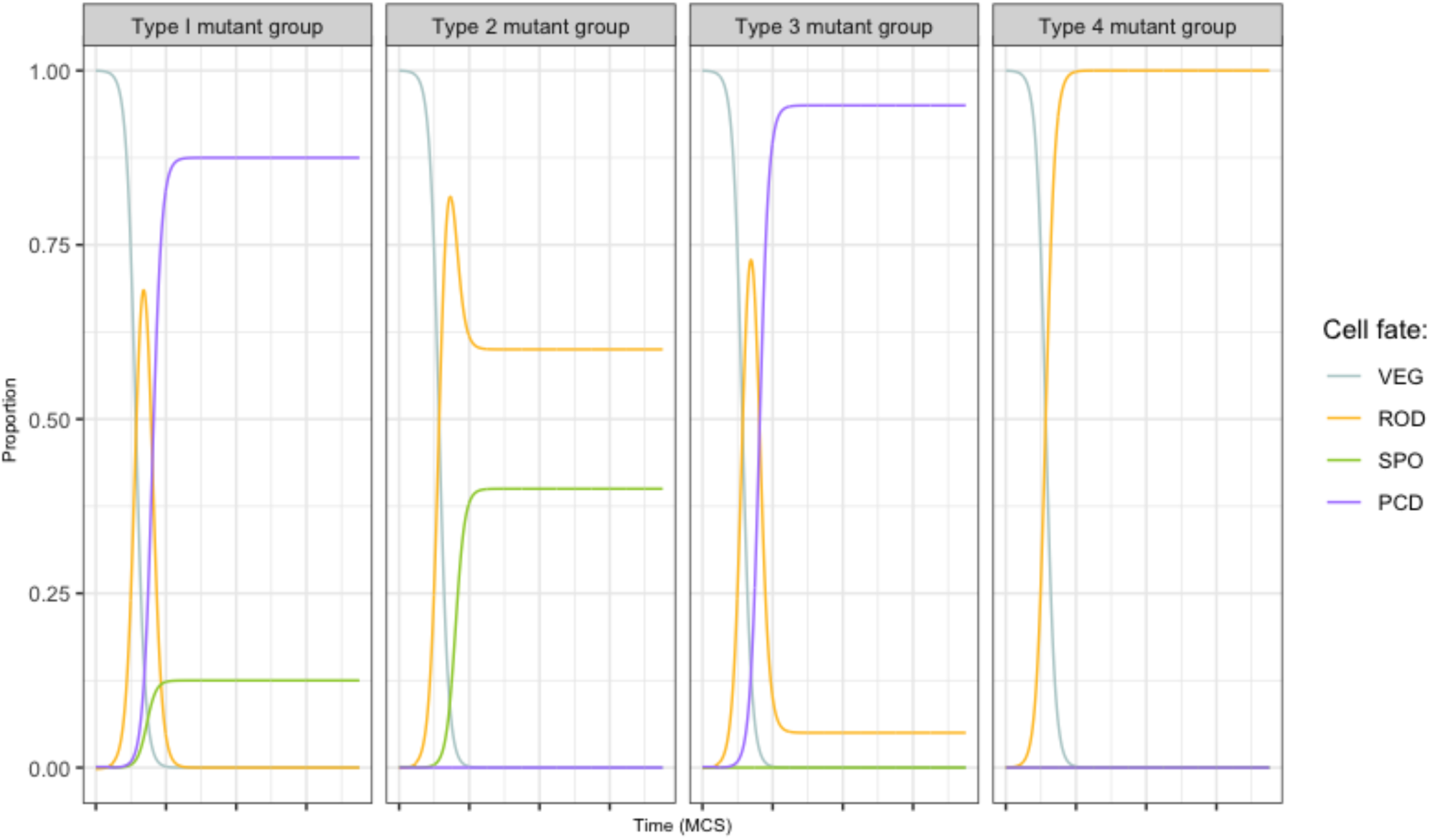
Effect of *in silico* knock-out mutants on cell fate trajectories. In each panel, the change of cell fate proportion over time is shown for a different mutant group. A mutant group comprises variables in the model (gene or proteins) that exhibit similar behaviour. No effect (Type 1), depletion of PCD (Type 2), depletion of SPO (Type 3), depletion of both PCD and SPO (Type 4). For variables contained in each mutant group see **Table S3**.

**Figure S4.**
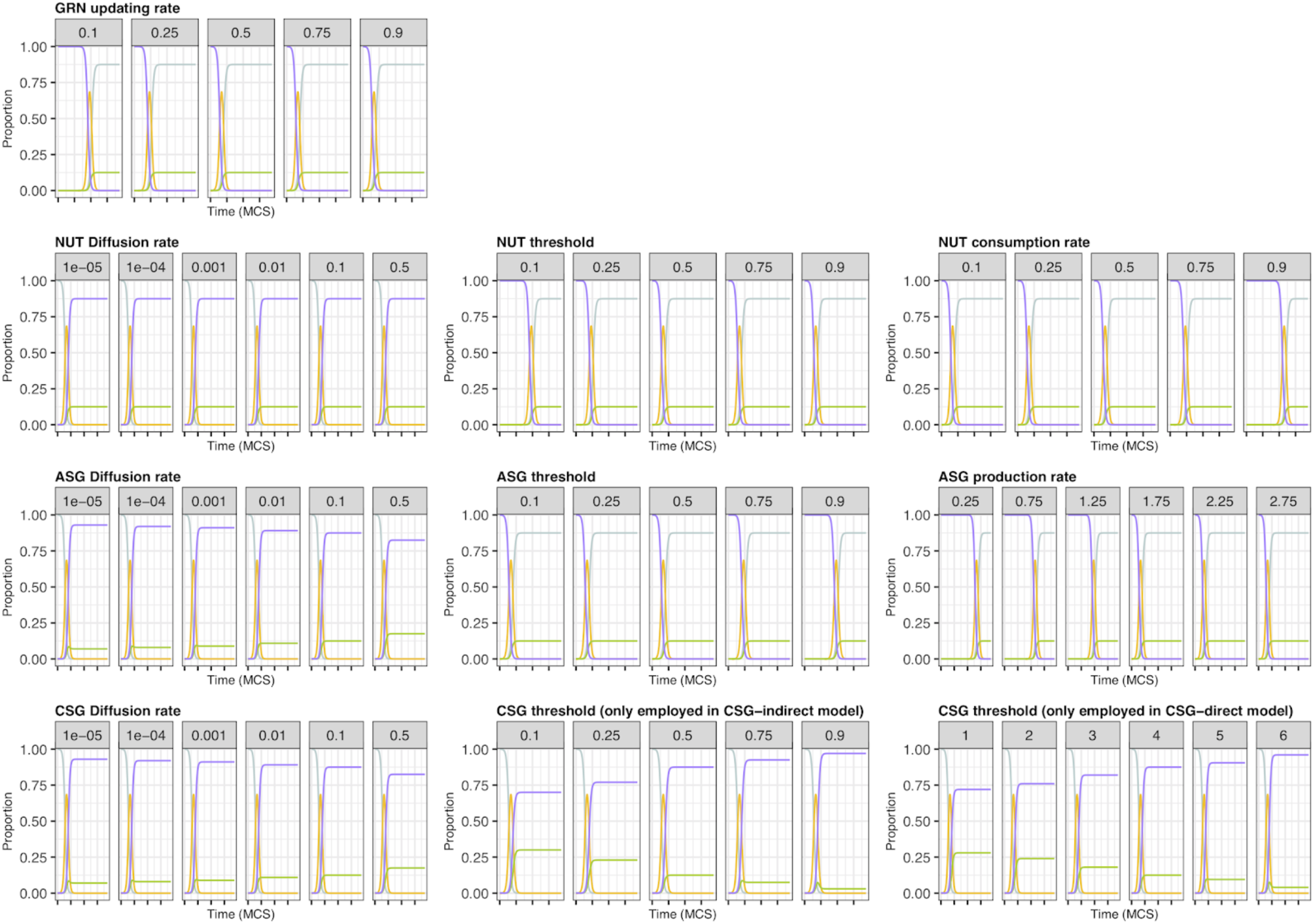
Sensitivity analysis for cell fate trajectories to parameter variation over time. Each panel shows cell fate trajectories over time upon variation in a single parameter. The modified parameter is annotated at the top of each panel (bold) and the specific value for each set of realization are shown in each sub-panel (gray boxes). Colored lines represent the proportion of individuals with each cell fates over time. Cell fates are VEG (cyan), ROD (yellow), SPO (green) and PCD (purple).

**Figure S5.**
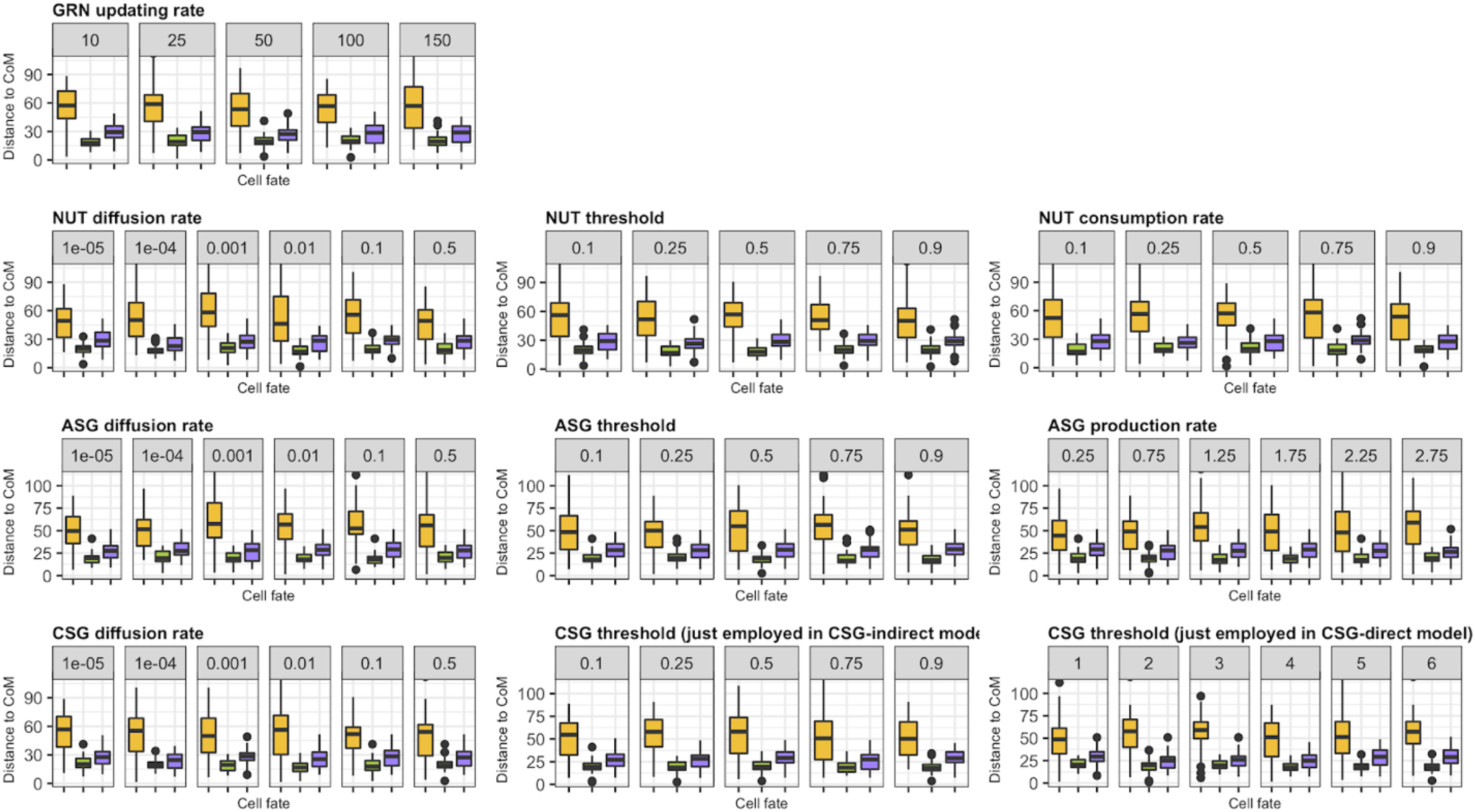
Sensitivity analysis of cell fate patterning to parameter variation. Each panel shows the distribution of distance relative to center of mass (CoM) for each cell fate upon variation in a single parameter. The modified parameter is annotated at the top of each panel (bold) and the specific value for each set of realizations are shown in each sub-panel (gray boxes). Boxplots represent the distribution of distance relative to center of mass for individual cell fates. Cell fates are ROD (yellow), SPO (green) and PCD (purple).

## 11. Supplementary Information 1

Specifications for the hybrid Boolean/ODEs GRN model.

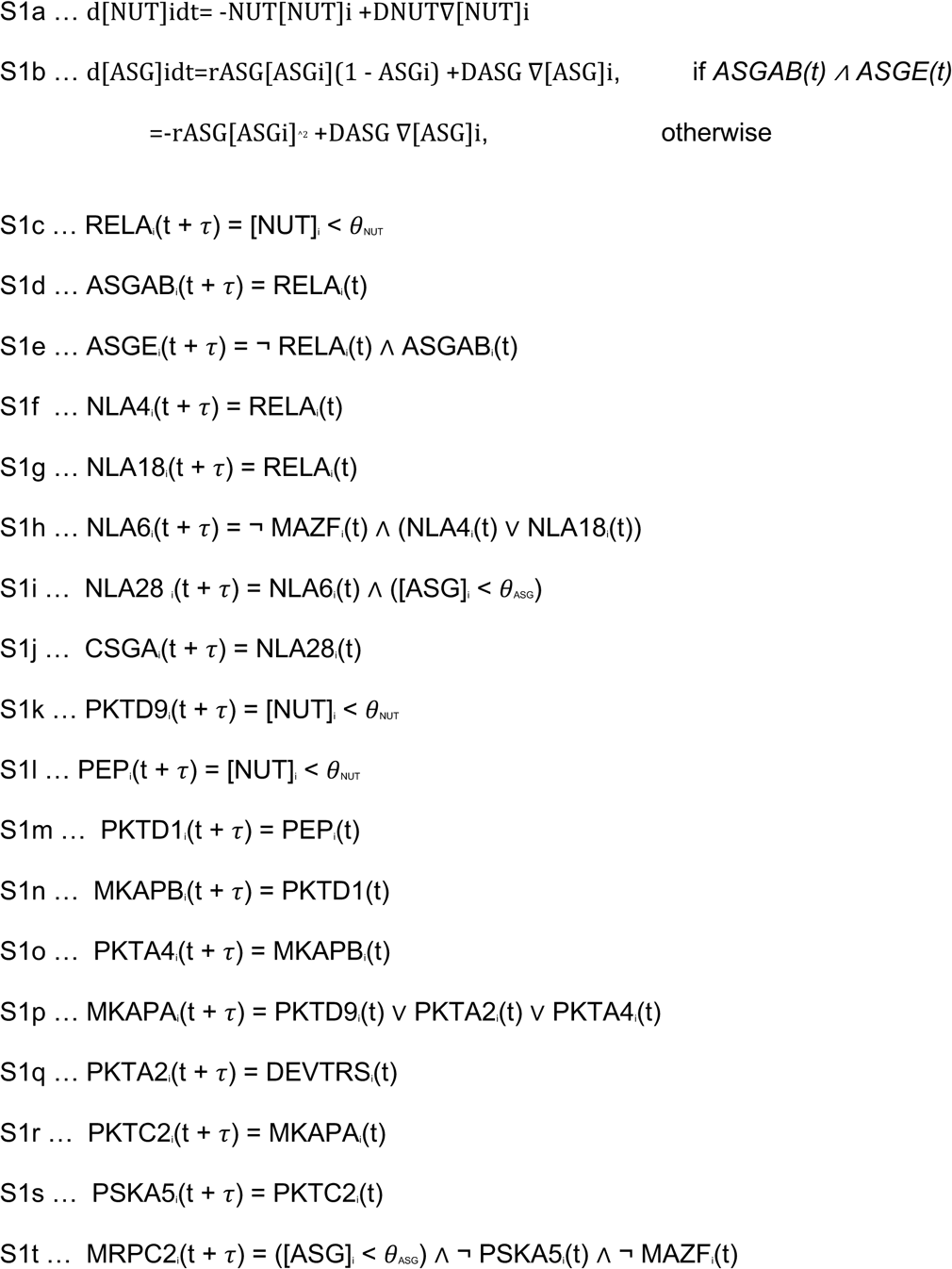

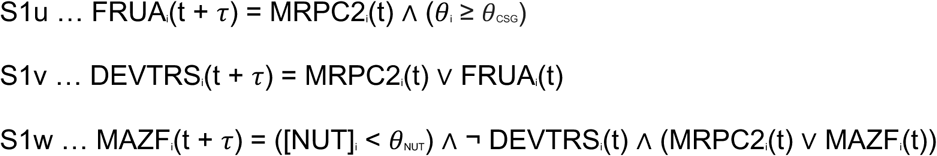

In equation S1u, *θ*_i_ is the number of neighbors to cell *i* with CSGAj(t) = 1.

## Supplementary Tables

**Table S1.** Model parameters. All parameters are measured in arbitrary units unless stated otherwise.

| Symbol | Description | Nominal value | Values tested | Eq. |
| --- | --- | --- | --- | --- |
| Potts models - *Cellular level* |  |  |  |  |
| $J(0,1)$ | Cell-to-medium adhesion energy | 12.0 | - | 1 |
| $J(1,1)$ | Cell-to-cell adhesion energy | 7.0 | - | 1 |
| $V_0(\sigma)$ | Target volume | 25.0 | - | 1 |
| $\lambda$ | Spring constant | 2.0 | - | 1 |
| T | Temperature | 10.0 | - | 2 |
| Boolean/ODEs GRN model - *Sub-cellular level* |  |  |  |  |
| $\tau$ | GRN updating rate | 50 MCS | 5, 10, 25, 50, 100 | S1a-u |
| $r_{\text{ASG}}$ | A-signal production rate | 1.75 | 0.1, 0.25, 0.5, 1.0, | S1b |
| $\gamma_{\text{NUT}}$ | Nutrient consumption rate | 0.5 | 0.1, 0.25, 0.75, 0.9 | S1a |
| $\theta_{\text{NUT}}$ | Nutrient threshold for starvation | 0.1 | 0.2, 0.25, 0.5, 0.75 | S1c,k,l |
| $\theta_{\text{ASG}}$ | A-signal activation threshold | 0.5 | 0.1, 0.25, 0.75, 0.9 | S1i,t |
| $\theta_{\text{CSG}}$ | Active neighbors threshold | 5 cells | 1, 2, ..., 6 | S1u |

| Partial differential equations - *Chemical field level* |  |  |  |  |
| --- | --- | --- | --- | --- |
| $D_{\text{ASG}}$ | A-signal diffusion rate | 0.5 | $10^{-5}$, $10^{-4}$, ..., $10^{-2}$, 0.1 | S1b |
| $D_{\text{NUT}}$ | Nutrient diffusion rate | 0.1 | $10^{-5}$, $10^{-4}$, ..., $10^{-2}$, 0.5 | S1a |
| $\rho$ | Nutrients released by dead cells | 10.0 | 0, 1, 5, 20, 100 | 3 |

**Table S2.** Summary of key parameters and their effects on properties in the model: ‘Yes’ means that the given property is sensitive to the parameter variation and ‘No’ that it is not.

| Parameter | Property |  |  |
| --- | --- | --- | --- |
|  | Temporality | Cell fate proportion | Cell fate patterning |
| *GRN updating rate* | No | Yes | No |
| *ASG diffusion rate* | No | Yes | No |
| *NUT diffusion rate* | No | Yes | No |
| *CSG diffusion rate* | No | Yes | No |
| *CSG threshold (direct model)* | No | Yes | No |
| *CSG threshold (indirect model)* | No | Yes | No |
| *ASG production rate* | Yes | No | No |
| *NUT consumption rate* | Yes | No | No |
| *NUT threshold* | Yes | No | No |
| *ASG threshold* | Yes | No | No |

**Table S3.** Classification of nodes included in the gene regulatory network by their ‘knock-out’ phenotypic effects.

| **Mutant group** | **Phenotype** | **‘Knock-out’ nodes** |
| --- | --- | --- |
| **Type 1** | Wild-type | PktA2, PktA4, PktD1, PskA5, PktC2, MkapA, MkapB |
| **Type 2** | No PCD | MazF |
| **Type 3** | No SPO | DevTRS, FruA, Nla4, Nla18, Nla6, Nla28, CsgA |
| **Type 4** | Neither PCD nor SPO | AsgAB, AsgE, MrpĆ2 |

